# Averaging in gene perturbation benchmarks hides perturbation-strength variation: Consequences for model evaluation

**DOI:** 10.64898/2026.10.20.760537

**Authors:** Yash Sharma, Madhav Subramaniyam, Pio Passariello, Dylan Jay, David Render

## Abstract

Perturbation-response benchmarks usually represent each genetic intervention by averaging all cells assigned that perturbation label. We inquire what perturbation-strength information is lost by this averaging, and whether current model predictions preserve enough perturbation-specific information for this perturbation-strength variation to be evaluated. Using X-Atlas/Orion CRISPRi screens, we use depth-corrected guide abundance as a cell-level proxy for perturbation strength and examine how transcriptional responses vary within a nominal perturbation. Higher perturbation strength is associated with greater response reproducibility. This effect is present in two human cell lines, HCT116 and HEK293T, but is larger in HCT116. For model evaluation, HCT116 retrieval tests show that observed responses preserve perturbation identity, whereas the evaluated STATE and Cell2Sentence predictions do not reliably recover it, hence the question of within-perturbation strength information cannot be meaningfully evaluated.

## 1 Introduction

Perturbation-response benchmarks usually assign one response target to each nominal intervention by averaging all cells carrying that label. This aggregation is often treated as a neutral way to reduce cell-level noise, but it also defines what the model is asked to predict. If biologically relevant response structure varies within the label, the pooled target marginalizes that structure rather than merely denoising it.

CRISPRi provides a concrete case [1]. It does not silence its target equally in every cell, so cells carrying the same nominal perturbation can differ in perturbation strength at the single-cell level. If the response varies with cell-level perturbation strength, two datasets can contain the same nominal perturbation yet define different prediction targets simply because their distributions of perturbation strength differ. Conventional perturbation-level evaluation cannot expose this dependence because cell-level perturbation strength is usually unobserved.

The X-Atlas/Orion CRISPRi screens make this hidden axis observable by providing per-cell guide abundance alongside gene expression [2]. This lets us ask two benchmark questions. First, when cells assigned the same perturbation have different cell-level perturbation strengths, what information is lost when those cells are collapsed into one perturbation-level target? Second, do current model predictions preserve enough perturbation-specific information for that finer within-perturbation variation to be evaluated?

For the first question, if the response is

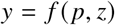

where *p* is perturbation identity and *z* is cell-level perturbation strength, a conventional perturbation-level benchmark instead uses

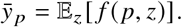

The benchmark therefore keeps which perturbation occurred but averages over cell-level perturbation strength.

Across HCT116 and HEK293T, we find a reproducible gene-expression change with perturbation strength that is shared across perturbations. This can make low- and high-strength groups appear to have different response directions even when the difference is not specific to the targeted gene. After estimating this shared component from negative-control perturbations and removing it, the apparent direction change disappears in both cell lines. Two independent samples of high-strength cells nevertheless give more similar estimates of the perturbation’s gene-expression effect than two independent samples of low-strength cells. This difference is large in HCT116 and smaller but statistically supported in HEK293T.

The second question concerns what information model predictions preserve. Perturbation strength is a finer distinction than perturbation identity. Before asking whether a prediction distinguishes between low and high strength cells within the same perturbation, it must first preserve which perturbation is being represented. In HCT116, independent observed responses recover perturbation identity, whereas the evaluated STATE predictions do not reliably exceed chance within its supported perturbation subset and Cell2Sentence predictions are essentially at chance. The evaluated zero-shot predictions therefore fail the perturbation-identity prerequisite required to test the finer within-perturbation strength axis. A perturbation-informed PCA reference illustrates the complementary limitation, that it preserves perturbation identity but uses the same strength-averaged template for both strength groups.

## 2 Related Work

Prior work has shown that cells carrying the same perturbation label need not have the same perturbation strength. Mixscape identifies cells that receive a perturbation label but show little detectable perturbation effect [3], while at-tenuated CRISPR guides have been used to deliberately titrate target-gene expression and produce graded downstream responses [4]. More recent methods infer continuous within-perturbation variation from the transcriptome itself. Mixs-cale estimates single-cell perturbation strength, while the perturbation-response score (PS) quantifies heterogeneous single-cell perturbation outcomes and enables dosage analysis without experimentally titrating the perturbation [5, 6]. X-Atlas/Orion itself uses sgRNA abundance as a proxy for knockdown efficacy, showing that low- and high-abundance strata differ in target-gene knockdown and, for essential-gene perturbations, in the magnitude of downstream transcriptional effects [2]. Because transcriptome-derived strength quantities are inferred from the same transcriptome whose response is later analyzed, the stratifying quantity and measured response are not fully independent; Mixscale uses leave-one-out procedures to reduce this dependence [5]. Guide abundance instead provides a measurement from the perturbation-delivery channel, separate from the gene-expression response being analyzed. We use depth-adjusted guide abundance to ask what information is lost when cells spanning this within-perturbation variation are averaged into a single benchmark target.

Perturbation-response datasets and prediction benchmarks commonly summarize cells at the nominal perturbation level, so predictions are compared with population-average responses [7, 8, 9, 10]. Recent work has shown that reported performance can depend strongly on baseline choice, reference construction, and expression patterns shared across perturbations rather than only on perturbation-specific prediction [11, 12, 13]. Population-denoising approaches address a different problem by estimating the perturbation-level average more accurately [14]. X-Cell and related evaluations likewise aggregate cells at the nominal perturbation level [15]. We ask instead what within-perturbation information is lost by that aggregation and whether model predictions preserve enough perturbation-specific information for that variation to be evaluated.

## 3 Methods

### 3.1 Problem setup and evaluation data

#### Perturbation-level benchmark target

Let *p* denote a nominal perturbation, *z* its cell-level perturbation strength, and *Y* the measured transcriptional response. Standard benchmarks aggregate all cells assigned to *p*, so their mean evaluation target is

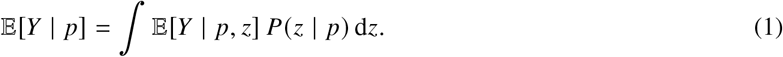

If the response changes with *z*, the pooled target depends on the distribution of perturbation strength *P*(*z* | *p*).

#### Datasets and perturbation-strength proxy

We use the X-Atlas/Orion HCT116 genome-scale dual-guide CRISPRi screen (3,409,169 cells × 38,606 genes) [2] as the primary context, and HEK293T from the same resource as a second cell line. Orion provides per-cell guide calls alongside expression, following direct guide-RNA capture in pooled single-cell screens [3]. For cell *i*, let *g*_*i*_ denote the smaller of its two guide UMI counts, *c*_*i*_ its total sequencing count, and *p*_*i*_ its perturbation. Within each perturbation, using evaluation-half cells, we regress log (1 +*g*_*i*_)on log (1 +*c*_*i*_)and use the residual as the perturbation-strength proxy,

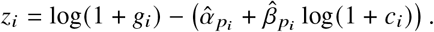

Evaluation-half cells are then divided within each perturbation into two quantile bins of *z*_*i*_, defining the lower and higher-strength groups. Each strength group is subsequently split into two disjoint halves for response estimation.

#### Held-out evaluation construction

Within each perturbation, cells are split into a selection half and an evaluation half. Analysis choices are fixed using the selection half, and all reported scores are computed on held-out evaluation cells. Perturbations with little measurable response on the selection half form a near-null group used as a negative control; they are not assumed to be biologically inert. A separate frozen perturbation set is used for model evaluation. Screening and split construction are detailed in Supplementary Sec. A.1.

### 3.2 Strength-resolved response analysis

To ask whether perturbation strength changes the perturbation response, we divide cells within each perturbation into lower and higher-strength groups. For each group, we split the cells into two independent samples and estimate the gene-expression response from each. Each sample is compared with a separate control population matched for sequencing depth and mitochondrial fraction. The correlation between the two response estimates measures how consistently independent cell samples recover the same perturbation response:

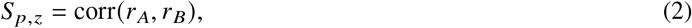

where *z* denotes the strength group. Full response estimation and matched-control construction are given in Supplementary Sec. A.2.

Cells with lower and higher perturbation strength can also differ in background properties. We estimate this common high-minus-low pattern from perturbations with little detectable response—near-null perturbations in HCT116 and null perturbations in HEK293T—and remove it using the direction estimated from the opposite split half. We then recompute response reproducibility after this correction. Our primary within-perturbation statistic is

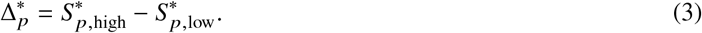

A positive value means that two independent samples of higher-strength cells give more similar estimates of the perturbation response than two independent samples of lower-strength cells. The full construction of the shared pattern and its removal is given in Supplementary Sec. A.2.

We separately ask whether the low-strength and high-strength responses differ in direction. Because noisy estimates of the same underlying response can appear less similar, we compare their observed similarity with the similarity expected from their measured split-half reproducibility. We treat this as a descriptive diagnostic and interpret it relative to negative-control and strength-shuffled comparisons. The full direction analysis is given in Supplementary Sec. A.2.

To test whether these differences genuinely depend on perturbation strength, we also shuffle strength labels among cells within each perturbation. This preserves the cells and perturbation identities while destroying the association between perturbation strength and transcriptional response. Confidence intervals are obtained by bootstrapping perturbations. Because the corrected statistic depends both on the perturbations being evaluated and on the negative controls used to estimate the shared high-minus-low pattern, both sets are resampled and the shared pattern is re-estimated in every bootstrap replicate. Full permutation and bootstrap procedures are given in Supplementary Sec. A.3.

### 3.3 Model evaluation

Let *p* denote perturbation identity and *z* ∈ {low, high} its strength group. The observed-data analysis asks whether the response depends on *z* within a fixed *p*. For a model, however, preservation of this finer information is only meaningful if its prediction first preserves *p* itself. We therefore test perturbation identity before asking whether predictions retain information about *z* within *p*. Moreover, the evaluated STATE and Cell2Sentence setups produce one prediction per nominal perturbation rather than separate predictions across *z*, which independently prevents a direct test of whether the predictions distinguish lower from higher strength.

Let *ŷ*_*p*_ denote the model-predicted response for perturbation *p*, and let 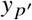 denote the observed response for candidate perturbation *p*^′^ ∈ P. For each candidate, we compute the response similarity

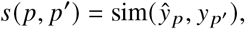

and rank all *p*^′^∈*P* by this score. If the model prediction preserves the characteristic response of perturbation *p*, the correct candidate *p*^′^ = *p* should rank near the top. Independent observed responses are evaluated in the same way as a positive control, showing how well perturbation identity can be recovered from the experimental data itself.

We summarize retrieval using mean reciprocal rank (MRR) and estimate chance performance by permuting perturbation identities. Candidate construction, gene filtering, similarity calculation, and permutation details are given in Supplementary Sec. A.4.

The PCA reference provides the complementary case. It learns a separate response template for each *p*, so perturbation identity is preserved, but the same template is used for both values of *z*. It therefore collapses within-perturbation strength variation by construction. Model-specific implementation details for PCA, STATE, and Cell2Sentence are given in Supplementary Sec. A.4.

## 4 Results

### 4.1 Perturbation strength varies measurably within a nominal perturbation

We first test depth-corrected guide abundance as a proxy for cell-level CRISPRi perturbation strength by asking whether higher values correspond to stronger suppression of the targeted gene. Across the 210 perturbations with a defined four-quartile correlation, the median Spearman correlation between mean strength proxy and relative target-gene expression across quartiles is −0.500. Target knockdown also increases monotonically across strength-proxy quartiles (Figure 1B–C). Thus cells carrying the same perturbation label show measurable differences in cell-level perturbation strength.

**Figure 1:**
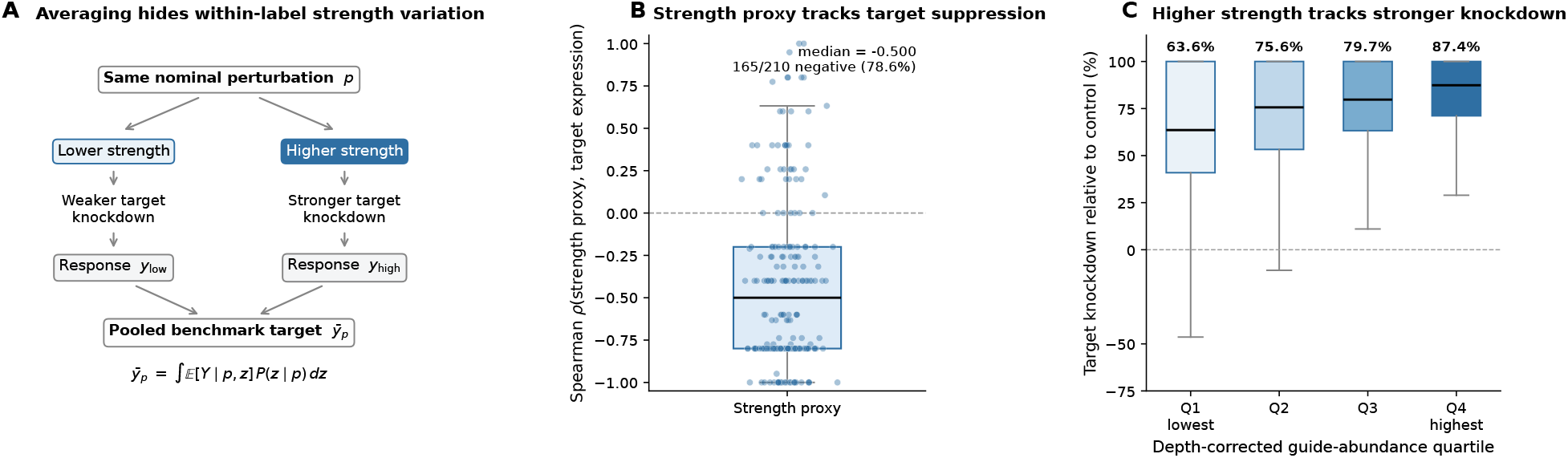
Perturbation strength varies within a nominal perturbation and tracks target-gene suppression. **(A)** Cells assigned the same nominal perturbation *p* can experience different perturbation strengths and therefore different degrees of target knockdown, producing distinct responses *y*_low_ and *y*_high_ that a perturbation-level benchmark pools into the single target 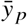 of Eq. (1). **(B)** Within-perturbation Spearman correlations between depth-corrected guide abundance and target-gene expression, across the 210 perturbations with defined quartile-level correlations; points show individual perturbations. **(C)** Target knockdown relative to control across the four strength-proxy quartiles, for the 252 perturbations contributing all four. Median knockdown per quartile is annotated above each box. In (B) and (C), boxes show the interquartile range, center lines the median, and whiskers extend to 1.5× the interquartile range.

Sequencing depth is an important alternative explanation because cells with more total counts also tend to have more guide counts. Residualizing guide abundance against depth removes the monotonic depth trend while strengthening the median correlation between guide abundance and target-gene expression from −0.400 to −0.500. We therefore use depth-corrected guide abundance to order cells by perturbation strength in the analyses that follow. HEK293T shows the same relationship, providing an independent validation in a second cell line (Sec. 4.4, Supplementary Sec. A.3). Because guide abundance is measured rather than experimentally assigned, these analyses establish association with the strength proxy rather than the causal effect of increasing perturbation strength.

### 4.2 Global controls can create apparent strength-dependent responses

Low and high strength cells can differ in background properties as well as perturbation strength. To test how much this can distort a response estimate, we perform a sham stratification in effect-free perturbations using mitochondrial function and compare both groups with the same global control population. Although this stratification is unrelated to perturbation strength, it produces apparent perturbation-specific signal as large as 0.480.

We therefore use separately matched control populations for each strength group. Even after this correction, however, low and high-strength cells can differ along an expression pattern shared across perturbations. The next analysis asks what perturbation-specific structure remains after accounting for this shared pattern.

### 4.3 In HCT116, higher perturbation strength improves response reproducibility after removing shared structure

Before removing the shared low-to-high expression pattern, strong HCT116 perturbations appear to show both increased response reproducibility and a change in response direction from low to high perturbation strength. The apparent direction change, however, is also present in medium, weak, and near-null perturbations, arguing against an effect specific to the targeted perturbation.

Independent estimates of the shared high-minus-low pattern from the two cell halves are strongly aligned under the original strength labels (cosine 0.789) but much less aligned after shuffling the labels (0.154; Figure 2A). After removing this shared component, the apparent direction change is reduced to approximately zero for both strong and near-null perturbations (Figure 2B). We therefore find no evidence of an additional perturbation-specific change in response direction with increasing perturbation strength.

**Figure 2:**
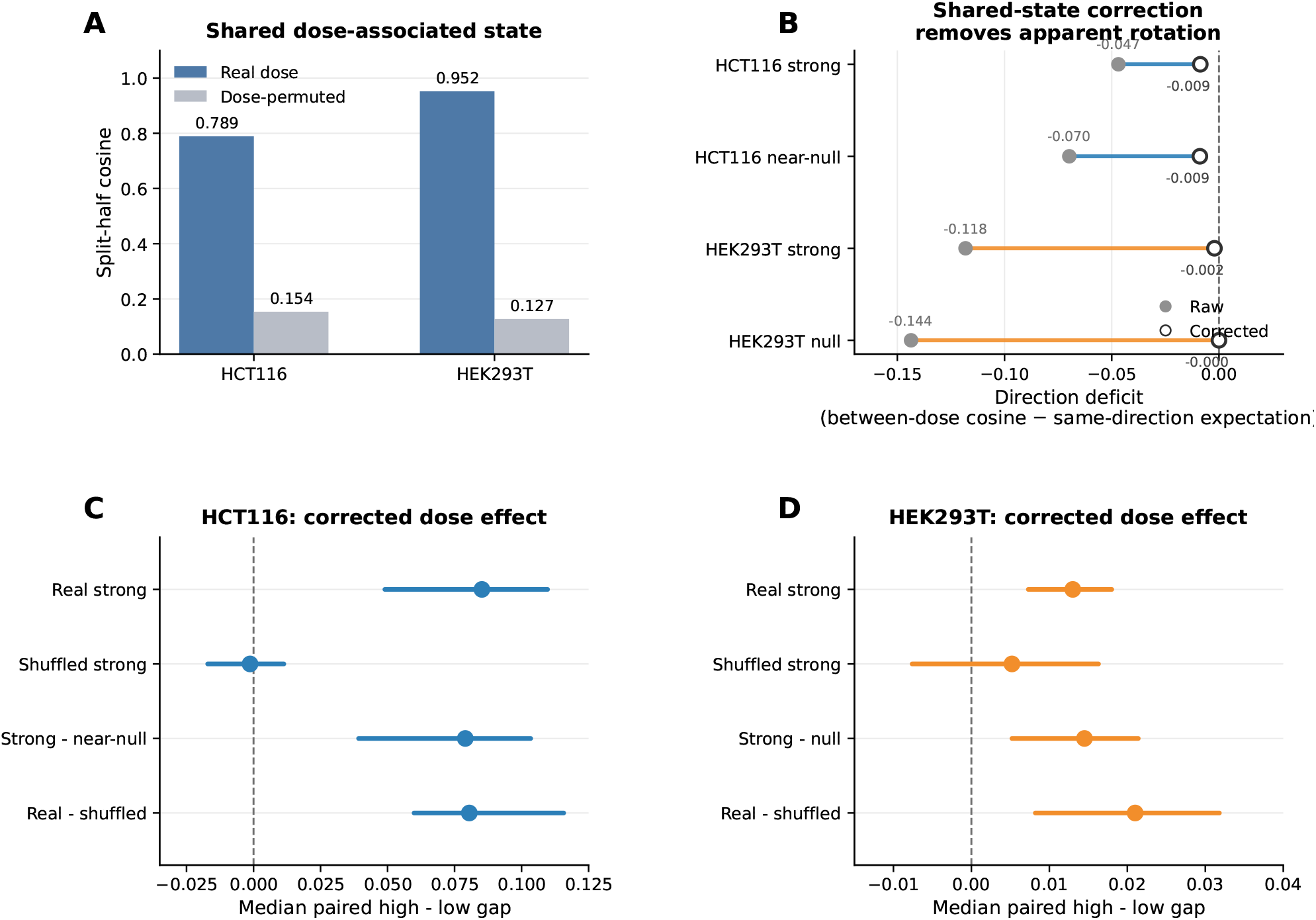
A shared strength-associated expression pattern explains the apparent change in response direction, while a perturbation-specific increase in response reproducibility remains. **(A)** Split-half cosine between the two independently estimated negative-control high-minus-low directions, using the original strength labels and after within-perturbation strength-label permutation, in each cell line. **(B)** Measured low-to-high cosine minus the cosine expected if both strength groups share an underlying response direction, accounting for their split-half response reproducibility. Filled markers show values before removing the shared pattern and open markers show corrected values; the difference approaches zero for strong and negative-control perturbations alike. **(C)** HCT116 median paired high-minus-low gaps with 95% nested bootstrap intervals, for the real strong stratum, the strength-shuffled strong stratum, and the strong-minus-near-null and real-minus-shuffled contrasts. **(D)** The same four quantities in HEK293T, with its null stratum in place of near-null. Note the different horizontal scales in (C) and (D).

The reproducibility increase survives the same correction. Two independent samples of high-strength cells give more similar estimates of the perturbation response than two independent samples of low-strength cells, while nearnull perturbations show little corresponding difference. For strong perturbations, the median paired high-minus-low reproducibility gap is +0.085 (95% bootstrap CI 0.049–0.110), whereas the strength-shuffled comparison remains centered near zero (Figure 2C). The strong-minus-near-null gap is +0.079 (95% CI 0.039–0.103), and the paired real-minus-shuffled strong contrast is +0.081 (95% CI 0.060–0.116). Thus the higher-strength response is more consistently recovered from independent cell samples, and this difference is not explained by the expression pattern shared across perturbations or by splitting cells according to perturbation strength.

### 4.4 HEK293T replicates a smaller increase in response reproducibility

HEK293T tests whether the same separation holds in another cell line. As in HCT116, low and high-strength cells initially appear to differ both in response reproducibility and response direction, including among null perturbations.

The shared high-minus-low expression pattern is particularly strong in HEK293T: independent estimates from the two cell halves have cosine 0.952 under the original strength labels but only 0.127 after shuffling (Figure 2A). Removing this pattern again eliminates most of the apparent change in response direction for strong and null perturbations alike (Figure 2B). The large uncorrected direction change is therefore dominated by expression changes shared across perturbations.

A smaller perturbation-specific reproducibility effect remains. The median paired high-minus-low gap for strong perturbations is +0.013 (95% bootstrap CI 0.007–0.018), while the shuffled comparison has an interval spanning zero (Figure 2D). HEK293T therefore reproduces the same qualitative result as HCT116: independent samples of higher-strength cells give more consistent estimates of the perturbation response, although the effect is substantially smaller.

### 4.5 Model predictions must preserve perturbation identity before strength can be tested

The observed-data results show that two pieces of information can matter: which perturbation occurred and its cell-level perturbation strength. We would ultimately like to ask whether a model distinguishes between low and high strength cells within the same perturbation. Before that question is meaningful, however, its prediction must first distinguish that perturbation from other perturbations.

We test this prerequisite using perturbation-identity retrieval. For each predicted response, we compare it with the observed responses of all candidate perturbations and ask where the correct perturbation ranks. If the prediction contains perturbation-specific information, its matching observed response should rank near the top. Independent experimental responses provide a positive control, showing whether perturbation identity is recoverable from the data itself.

For Cell2Sentence, independent observed responses achieve an MRR of 0.165, whereas Cell2Sentence reaches only 0.010, indistinguishable from its permutation null (Figure 3A). Thus the experimental responses contain perturbation-specific information that is largely absent from the evaluated Cell2Sentence predictions.

**Figure 3:**
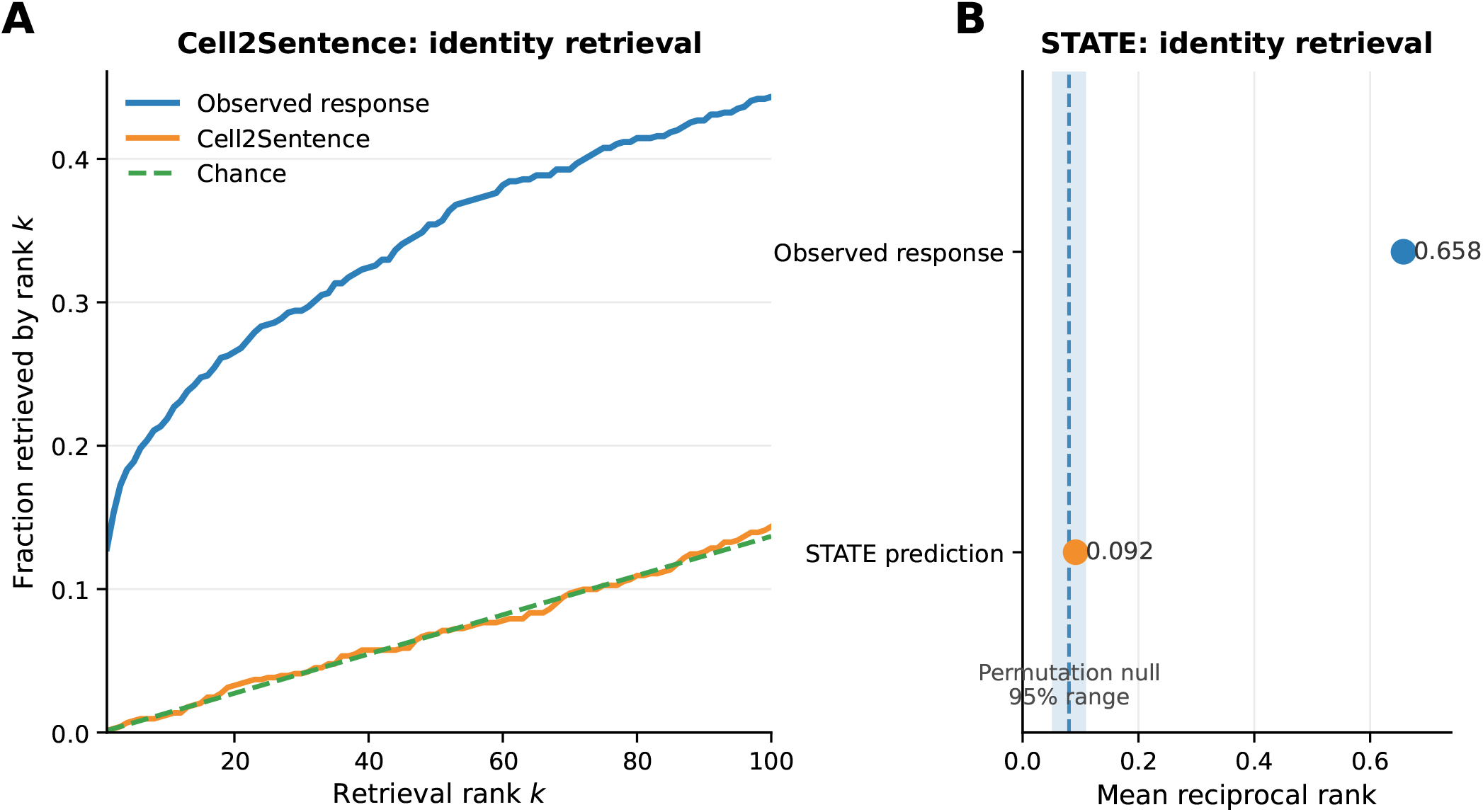
In HCT116, observed responses preserve perturbation identity that the evaluated zero-shot predictions fail to recover in their valid retrieval universes. **(A)** Cumulative retrieval over the 731-perturbation candidate set (756 downstream genes): the fraction of queries whose true perturbation is retrieved within rank *k*. Independent observed responses separate clearly from the plotted chance curve (MRR 0.165, *p*_perm_ = 0.0005), whereas Cell2Sentence tracks it (MRR 0.010, *p*_perm_ = 0.35). **(B)** Mean reciprocal rank over the 62 perturbations present in STATE’s perturbation mapping (774 downstream genes). Observed responses reach 0.658 (*p*_perm_ = 0.0005), while STATE predictions reach 0.092 (*p*_perm_ = 0.144), inside the shaded 95% range of the 2,000-permutation identity null. In both panels, perturbation target genes are excluded and the generic observed response direction is projected out before cosine retrieval.

STATE requires restricting the analysis to the 62 candidate perturbations supported by its released perturbation vocabulary. Within this valid subset, independent observed responses are strongly identifiable (MRR 0.658), whereas STATE predictions achieve an MRR of 0.092 and do not significantly exceed the permutation null (Figure 3B).

The evaluated Cell2Sentence and STATE predictions therefore fail the perturbation-identity prerequisite: unlike independent observed responses, they do not reliably preserve the coarser information about which perturbation occurred. The finer question of whether they preserve within-perturbation strength information consequently cannot be assessed from these predictions. This does not show that the models cannot encode strength in principle; it shows that their evaluated predictions do not preserve enough perturbation-specific information for that test.

PCA provides the complementary case. It learns a separate template for each perturbation, so perturbation identity is preserved, but that template is estimated after averaging across perturbation strength and then reused unchanged for lower and higher-strength cells. It therefore collapses the within-perturbation strength axis by construction.

## 5 Discussion

### Averaging over perturbation strength changes the benchmark target

Equation (1) makes explicit that a perturbation-level target depends not only on perturbation identity but also on the distribution of cell-level perturbation strengths among the measured cells. Two screens can therefore contain the same nominal intervention yet define different aggregate targets if their perturbation-strength distributions differ, even when the underlying strength-conditional responses are the same. Population-denoising methods can estimate that aggregate more accurately [4], but they cannot recover variation that has already been averaged over. Cell-level perturbation strength therefore adds a within-perturbation source of benchmark heterogeneity to the between-perturbation effects emphasized in previous work [5].

### The corrected effect concerns the component not shared across perturbations

The low-to-high expression change contains a reproducible component shared across perturbations. After removing this component, higher perturbation strength is still associated with greater split-half response reproducibility in both cell lines, whereas the apparent perturbation-specific change in response direction largely disappears. The interpretation of this correction is that projection also removes any genuine perturbation-specific response that happens to align with the shared direction. The corrected statistic therefore describes the response component orthogonal to that shared pattern, not the complete biological effect of perturbation strength. Nor does the analysis determine whether the shared pattern itself reflects technical variation, biological cell state, or a combination of the two. More generally, strength-resolved evaluation requires reference populations and negative controls that distinguish perturbation-specific effects from background structure associated with the stratification itself [6].

### Strength-aware model evaluation has a perturbation-identity prerequisite

Evaluating whether a prediction preserves within-perturbation strength information is a finer question than evaluating whether it preserves perturbation identity. The HCT116 retrieval experiments show that perturbation identity is recoverable from independent observed responses, but is not reliably recovered by the evaluated Cell2Sentence or vocabulary-supported STATE predictions. Their within-perturbation strength representation therefore cannot be meaningfully assessed with this test. The PCA reference illustrates the complementary limitation: perturbation identity is preserved, but within-perturbation strength variation is collapsed by construction. These results do not establish that either zero-shot architecture is incapable of representing strength; they show that the evaluated outputs do not satisfy the prerequisite needed to test it. Perturbation-identity retrieval is also distinct from diagnostics such as gene-coordinate permutation [7]: each tests a different form of information preservation.

### Scope and limitations

Depth-corrected guide abundance provides a proxy for cell-level perturbation strength rather than a direct measurement or an experimentally assigned perturbation level, so its association with transcriptional response should not be interpreted as the causal effect of increasing perturbation strength. Residualizing sequencing depth and matching controls reduce specific sources of confounding but cannot eliminate all possible differences between lower and higher-strength cells. A gain in split-half reproducibility also does not imply a larger response magnitude; it means that independent samples recover the response more consistently. The corrected effect is substantially smaller in HEK293T than in HCT116, indicating that its magnitude is context dependent. We use two strength groups because a three-bin analysis introduces structure in the negative controls (Supplementary Sec. A.2). Finally, full perturbation-identity retrieval is evaluated only in HCT116; the HEK293T Cell2Sentence null-token ablation is not a matched cross-cell-line retrieval test.

## 6 Conclusion

A nominal perturbation does not by itself determine the target used by a perturbation benchmark. When cells with different perturbation strengths are averaged, the target also depends on the distribution of strengths in the measured population. Our results show that this within-label variation is not simply removable noise. Even after accounting for expression changes shared across perturbations, responses estimated from higher-strength cells are more reproducible. Aggregation can therefore discard structured information before a predictive model is ever evaluated.

This separates two sources of information loss that conventional benchmark scores can conflate. The benchmark can remove within-perturbation variation when it constructs a single target, and a predictive model can fail to preserve the perturbation-specific response represented by that target. Strength-aware evaluation should therefore distinguish information removed by benchmark construction from information lost in model prediction.

## Acknowledgements

We would like to thank Dr Katalin Susztak and Chenyu Li from Perelman School of Medicine at the University of Pennsylvania for the helpful discussions.

## A Supplementary Material

### A.1 Data and screening

#### Depth residualization

Raw guide abundance correlates with sequencing depth (*ρ* = 0.249). Median sequencing depth increases across raw guide-abundance quartiles (13,994, 15,947, 17,992, 22,128 counts). We residualize log (1+ *g*_*i*_)against log (1 +*c*_*i*)_ within each perturbation, where *g*_*i*_ is guide abundance and *c*_*i*_ is total sequencing count. For the strength-resolved response analysis, this regression is fit separately within each perturbation using its evaluation-half cells, and the resulting residual defines the cell-level perturbation-strength proxy *z*_*i*_. The depth medians across depth-corrected guide-abundance quartiles are no longer monotonic (17,282, 16,471, 17,114, 18,225). The median Spearman correlation between the strength proxy and target-gene expression changes from −0.400 to −0.500, and the fraction of perturbations with decreasing target expression from 69.5% to 78.6%.

#### Perturbation-set construction

We construct two frozen perturbation sets using selection-half measurements, with downstream evaluation performed on held-out cells (Figure 4).

**Figure 4:**
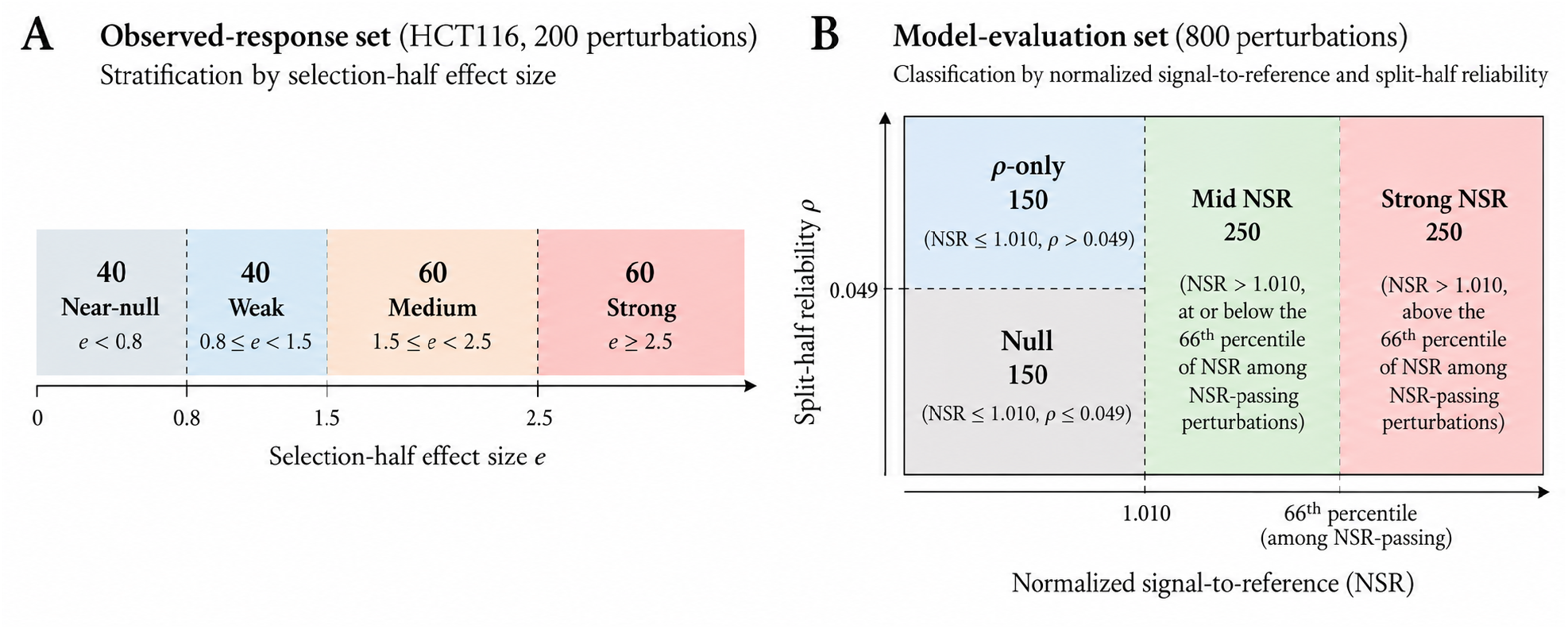
Construction of the two frozen perturbation sets. (A) HCT116 observed-response perturbations stratified by selection-half effect size. (B) Model-evaluation perturbations classified by normalized signal-to-reference (NSR) and split-half reliability (*ρ*). Numbers indicate the selected perturbations in each group. Regions illustrate classification rules rather than the observed distribution of perturbations.

#### Observed-response set

The HCT116 observed-response analysis uses 200 perturbations stratified by the magnitude of their transcriptional response, estimated from selection-half cells. The near-null group serves as a negative control.

#### Model-evaluation set

The model experiments use a separate set of 800 perturbations screened on selection-half cells by normalized signal-to-reference (NSR) and split-half reliability *ρ*. For perturbation *p*,

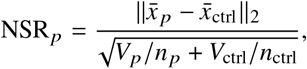

where *V* is the sum of per-gene variances, while *ρ* _*p*_ is the across-gene correlation between responses estimated from two random halves of the perturbation’s cells. Thresholds are the 95th percentiles of 300 size-matched relabelled-control draws. After requiring at least 60 evaluation-half cells, we randomly sample 250 strong-NSR, 250 mid-NSR, 150 *ρ*-only, and 150 null perturbations; strong- and mid-NSR are separated at the 66th percentile of NSR among retained perturbations.

### A.2 Response estimation

#### Matched controls and strength resolution

Each strength group is matched separately on a 10× 8 quantile grid of log-transformed sequencing depth and mitochondrial fraction (pct counts mt). For each perturbed split half, control means from the independent control half are weighted by the fractions of perturbed cells occupying each grid cell. Missing grid-cell control means are replaced by that control half’s pooled mean. The median weight requiring this fallback is zero in both cell lines.

Three strength bins produced a middle-bin value below the lowest bin and a strength gradient among effect-free controls. We therefore use two strength bins. Within each perturbation, evaluation-half cells are divided at the median of *z*_*i*_ into lower and higher-strength groups. Each group is then split into two disjoint halves *A* and *B* for response estimation. Perturbations for which two distinct strength groups cannot be formed are excluded. Strength-label permutations repeat the same matching and correction procedure.

#### Response vectors and corrections

For each perturbation *p* and strength group *z*, we split the perturbed cells into disjoint halves *A* and *B* and estimate the gene-expression response independently in each half. Each perturbed half is compared with its own matched control baseline. If *μ*_*h*_ is the mean count-fraction profile of perturbed cells in half *h* and *c*_*h*_ is its matched control profile, the log-fold-change response is

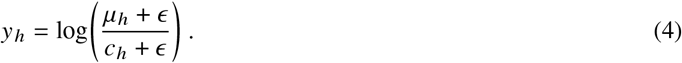

Before comparing responses across perturbations, we remove an expression-change direction that is common to many perturbations. Without this correction, two perturbations can appear similar simply because both contain the same generic perturbation-associated expression pattern. We estimate this common direction from a pooled profile *μ*_*U*_ containing up to 3,000 evaluation-half perturbed cells and compare it with the matched control baseline *c*_*U*,*h*_ in each control half:

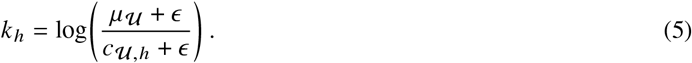

The perturbed-cell profile *μ*_*U*_ is shared between *k* _*A*_ and *k* _*B*_, while their matched control profiles are independently split. The pooled profile is constructed across perturbations without leaving out the perturbation being scored, so cells from that perturbation may also contribute to *μ*_*U*_.

With 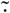 denoting gene-centering, we remove from each response the component aligned with the common direction estimated using the opposite control half:

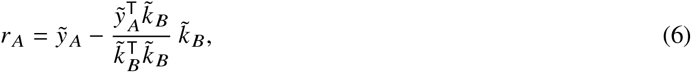

with the corresponding *r*_*B*_ using *k* _*A*_. Using the opposite half reduces dependence between the response estimate and the direction used to correct it.

The direction *k* above captures expression changes shared across perturbations in general. We separately ask whether moving from low to high perturbation strength produces an expression change that is itself shared across perturbations. We estimate this low-to-high pattern from negative-control perturbations: near-null perturbations in HCT116 and null perturbations in HEK293T.

For each negative-control perturbation *p*, its high-minus-low response in each half is

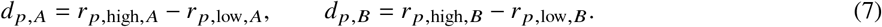

Averaging these differences across the negative-control set N gives the shared strength-associated directions

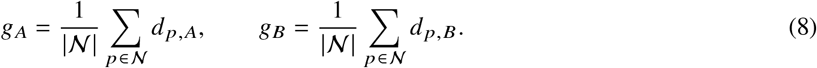

Thus, *k* represents expression structure shared across perturbations, whereas *g* represents the expression change shared specifically when moving from low to high perturbation strength.

We remove the component of each response aligned with this shared strength-associated direction, again using the direction estimated from the opposite half:

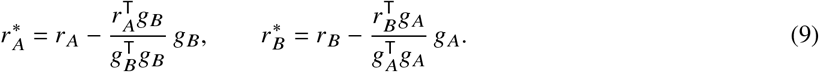

When a negative-control perturbation is itself being evaluated, its own high-minus-low vector is excluded from the corresponding estimate of *g*. This prevents that perturbation from helping define the shared direction against which its own response is corrected, leaving |*N* | −1 perturbations in each equal-weight average.

The corrected within-strength response reproducibility is

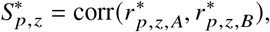

which measures how similarly two independent cell samples recover the corrected perturbation response. The primary high-minus-low reproducibility gap is defined in Eq. (3).

We separately test whether low- and high-strength responses differ in direction. A low similarity between their estimated response vectors does not by itself imply a biological direction change: noisy estimates of the same underlying response can also appear dissimilar. We therefore first measure the cross-half similarity between low- and high-strength responses,

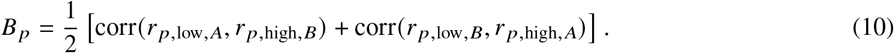

We then use the split-half reproducibility of the low- and high-strength responses to estimate how similar they should appear if both share the same underlying direction but are measured with different amounts of noise:

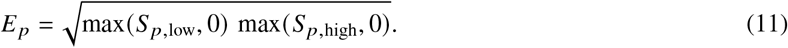

Nonfinite split-half correlations give a missing expectation, and negative values are clipped to zero. We use *B* _*p*_− *E* _*p*_ as a descriptive direction-change diagnostic and interpret it only relative to the negative-control stratum and strength-shuffled controls, before and after removing the shared strength-associated direction.

### A.3 Statistical inference

We use 20,000 perturbation-level bootstrap replicates. Each replicate independently resamples signal and negative-control perturbations with replacement. The resampled negative controls re-estimate both split-half shared strength-associated directions; the resampled signal perturbations determine the median corrected high-minus-low response reproducibility gap. Original-label and strength-shuffled comparisons use the same sampled signal perturbations.

Paired gaps are computed within perturbation before taking the median; they need not equal the difference between separate low- and high-strength medians. Table 3 gives the HCT116 bootstrap contrasts.

Secondary HCT116 tests give Wilcoxon *p* = 2.07 ×10^−9^ for the strong paired gap and *p* = 1.32 ×10^−8^ for real versus shuffled, and Mann–Whitney *p* = 5.26×10^−8^ for strong versus near-null gaps. For HEK293T, the corresponding conditional tests give *p* = 3.98×10^−9^, 1.27×10^−4^, and 8.18×10^−5^. The bootstrap intervals reported in the main text are primary because they include re-estimation of the shared strength-associated direction.

#### Additional HEK293T results

Median target knockdown is 51.5% in HEK293T and 75.4% in HCT116. Before shared-strength correction, HEK293T mid-NSR response reproducibility increases from 0.0831 to 0.1234 across strength bins, and *ρ*-only from 0.0875 to 0.1381; the corresponding strength-shuffled values are nearly flat. Corrected strong- and null-stratum results appear in Sec. 4.4 and Table 1.

**Table 1:** Cell-level perturbation strength is a reproducible within-label axis in both cell lines. We estimate the shared expression-change pattern from perturbations with little detectable response and subtract it using the opposite split half. After correction, higher perturbation strength is associated with increased perturbation-specific response reproducibility in both HCT116 and HEK293T, with a substantially larger effect in HCT116. Arrows show low and high-strength medians computed separately; paired gaps and real-minus-shuffled contrasts take within-perturbation differences before the median and therefore need not equal differences of displayed medians.

|  | HCT116 | HEK293T |
|---|---|---|
| Median strength-proxy–target correlation | −0.500 | −0.80 |
| Perturbations decreasing with strength proxy | 78.6% | 83% |
| Original-label shared-direction cosine | 0.789 | 0.952 |
| Shuffled shared-direction cosine | 0.154 | 0.127 |
| Raw strong, low → high | 0.152 → 0.232 | 0.108 → 0.138 |
| Raw negative-control, low → high | 0.026 → 0.070 | 0.074 → 0.106 |
| Corrected strong, low → high | 0.091 → 0.197 | 0.015 → 0.021 |
| Corrected negative-control, low → high | 0.004 → 0.007 | 0.011 → 0.011 |
| Corrected paired strong gap | +0.085 | +0.013 |
| Strong minus negative-control gap | +0.079 | +0.015 |
| Real minus shuffled strong gap | +0.081 | +0.021 |

**Table 2:** Information required for strength-aware evaluation and retained by each HCT116 model arm. Perturbation identity is assessed by retrieval within each arm’s valid candidate set (Table 4). Strength structure is only testable once identity is recovered; the PCA reference is the complementary case, preserving perturbation identity while averaging perturbation strength into a single perturbation-level template.

| Representation | Perturbation identity | Within-perturbation strength |
|---|---|---|
| Observed HCT116 response | Recoverable | Stronger at high strength |
| STATE (zero-shot, supported) | Near chance | Not testable |
| Cell2Sentence (zero-shot) | Near chance | Not testable |
| PCA reference | Preserved | Collapsed by construction |

**Table 3:** Primary inference for the HCT116 perturbation-strength effect after removing the near-null-derived shared strength-associated direction. The nuisance direction is re-estimated inside every bootstrap replicate.

| Corrected contrast | Estimate | 95% bootstrap CI |
|---|---|---|
| Real strong high-minus-low | +0.0852 | [0.0490, 0.1097] |
| Shuffled strong high-minus-low | −0.0013 | [−0.0171, 0.0113] |
| Real strong minus near-null gap | +0.0790 | [0.0392, 0.1034] |
| Real minus shuffled strong gap | +0.0805 | [0.0599, 0.1157] |

**Table 4:** Perturbation-identity retrieval using independent observed responses or model predictions as queries. Each query is ranked against the corresponding held-out response universe (|*P*| = 62 or 731). Chance median ranks are 31.5 and 366, respectively. STATE’s top-10 permutation *p* = 0.076.

| Query | \|𝒫\| | Top-1 | Top-10 | Median rank | MRR | *p*<sub>perm</sub> |
|---|---|---|---|---|---|---|
| *62-candidate evaluation* |  |  |  |  |  |  |
| Observed (independent half) | 62 | 36/62 | 49/62 | 1 | 0.6577 | 0.0005 |
| STATE | 62 | 2/62 | 14/62 | 32.5 | 0.0916 | 0.144 |
| *731-candidate evaluation* |  |  |  |  |  |  |
| Observed (independent half) | 731 | 94/731 | 160/731 | 137 | 0.1647 | 0.0005 |
| Cell2Sentence | 731 | 1/731 | 9/731 | 363 | 0.0100 | 0.35 |
| Cell2Sentence, null token | 731 | 0/731 | 10/731 | 366 | 0.0093 | 0.88 |

### A.4 Model implementation and retrieval details

#### Strength-blind PCA reference

Principal components are fit on control cells. A perturbation-specific shift is learned from selection-half cells in PC space and reused across both strength groups; it is a strength-blind reference, not a zero-shot model.

#### STATE

We evaluate the official ST-SE-Replogle RPE1 zero-shot checkpoint (best.ckpt) [8]. Its fixed perturbation mapping covers 485/ 5,882 HCT116 targets in the screen. Unsupported targets use the fallback control encoding, so their previous zero Overlap@*N* scores do not measure predictive performance.

#### Cell2Sentence

We evaluate *C2S-Scale-Pythia-1b* [9] zero-shot. Predicted gene ranks are converted to expression using a linear fit of log expression on log rank. On 800 frozen perturbations (19,200 cells), real versus null perturbation tokens change Pearson, Spearman, and cosine by at most 4 × 10^−4^.

#### Perturbation-identity retrieval

Of the 800 frozen HCT116 perturbations, 731 have targets in the common retrieval gene vocabulary. Cell2Sentence uses these 731 candidates and 756 downstream genes after control-expression filtering and removal of target genes. For STATE, only 62 of these candidates occur in the checkpoint’s perturbation mapping; we rerun retrieval and permutation tests in that 62-candidate universe. Its 1,957 overlapping output genes are filtered above the 60th control-expression percentile and stripped of perturbation targets, leaving 774 downstream genes. Before cosine retrieval, the generic observed-response direction is projected out of the compared profiles. Retrieval significance uses 2,000 identity-label permutations, with *p*_perm_ = (1+ *b*) /2001, where *b* is the number of permuted statistics at least as large as the observed statistic.

